# Alternative promoters used during myeloid differentiation and upon activation change the gene products available for innate immune programs

**DOI:** 10.64898/2026.02.01.703142.1

**Authors:** Miguel Ángel Berrocal-Rubio, Josie Gleeson, Masaki Kato, Diane Delobel, Hitesh Kore, Anthony G. Beckhouse, Dipti Vijayan, Kelly Hitchens, Takeya Kasukawa, Chi Wai Yip, Chen Zhan, Michael B Clark, Benjamin L. Parker, Hazuki Takahashi, Piero Carninci, Suzanne K Butcher, Christine A Wells

**Author notes:** COI JG has received support from Oxford Nanopore Technologies (ONT) to present their findings at scientific conferences. COI AGB is a currently employed by Illumina. COI DV is a currently employed by CSL.

## Abstract

Macrophages are innate immune cells present in most tissues of the body, whose molecular programs are determined by their ontogeny and environment. From the earliest stages of embryonic development, macrophages are recruited into developing tissues where they support organogenesis with trophic factors such as WNT, VEGF and PDGF. While macrophage subsets have been described in different tissues at single cell resolution, little is known about transcript isoforms and proteoforms that underpin their differentiation and function. Here we assessed enhancer, promoter, transcript and proteomic variation as pluripotent stem cells differentiate to macrophages, identifying over 200 previously uncharacterised genes and over 20,000 new mRNA isoforms, updating our current understanding of the human genome, its regulation and potential output. Newly discovered myeloid-expressed transcripts and proteins were enriched for motifs associated with secreted proteins, and these included previously uncharacterised isoforms of growth factors, in which we predict N-terminal changes impact on their location and function. Activation of primary adult monocytes and monocyte-derived macrophages was also characterised by the expression of diverse transcript isoforms, largely arising from alternate transcription initiation sites and predicted to impact on the acute response to bacterial or fungal stimuli. Understanding the full spectrum of gene products expressed by these cells further extends our understanding of the phenotypic plasticity and trophic potential of macrophages in human development and may lead to the discovery of new clinical targets for tissue engineering or immune-related studies.

**Graphical Abstract:** In this manuscript, Berrocal-Rubio and colleagues examined the differentiation of human macrophages using an iPSC model of tissue macrophage biology. Combining long-read sequencing technology with promoter profiling identified over 17, 700 genes implicated in pluripotency-myeloid specification. 7% of transcripts profiled from previously characterised genes were predicted to encode new proteins, and a further 3% of transcripts were derived from genes newly discovered in this project. They confirmed that a high proportion of these alternate transcripts were detectable in primary monocytes but also discovered that activation of primary monocytes led to further alternate promoter usage, with the potential to further diversify the innate immune responses to a broad set of pathogens. The newly described macrophage genes and transcripts encoded proteins enriched for motifs associated with secreted peptides. These data suggest that alternate transcription of macrophage genes leads to new effectors of innate immune function, that include a substantially expanded number of growth factors or secreted proteins. Created in BioRender. Berrocal, M. (2025) https://BioRender.com/d0n47le

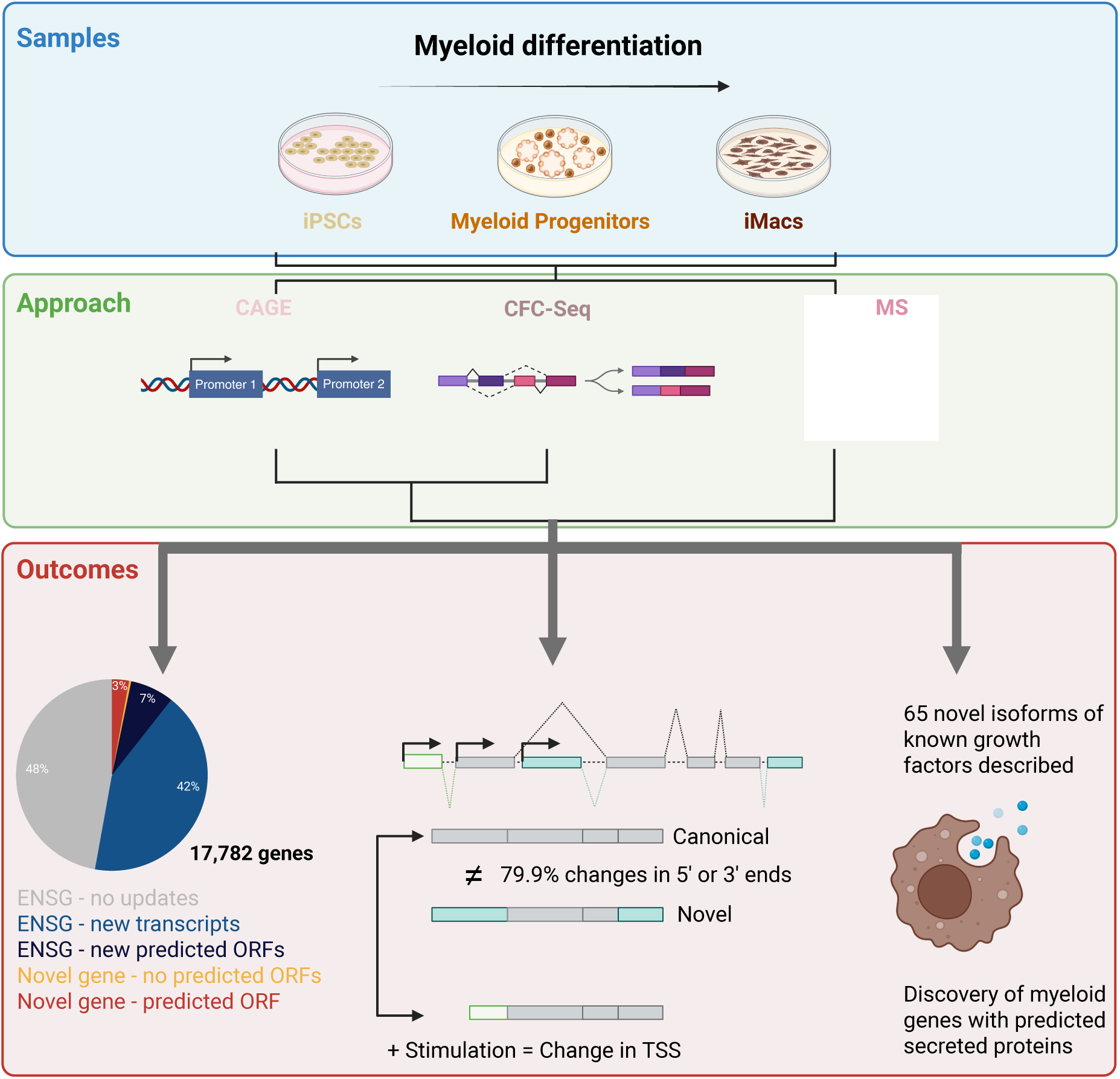

## Main

Macrophages are major regulators of homeostasis in virtually every human tissue. These cells are present from the earliest stages of human embryonic development (Carnegie Stage 7), where they have a morphogenic and reparative phenotype^1,2^. Maintenance of tissue homeostasis is a key function of macrophage populations throughout development to adulthood, and is characterised, among other things, by expression of a variety of growth factors^3^. Historically, studying the establishment of these early myeloid lineages has been challenging due to scarce data on early human embryos, although recent single cell studies in the developing human yolk sac and embryo have highlighted differences between mouse and human myelopoiesis^1^. In several animal models, macrophage-derived growth factors have been implicated in healthy mouse lymphatic development^4^, skin development^5^, and heart homeostasis^6^ but how macrophage-derived factors differ from other stromal signals is not well understood. Addressing this question requires the increased molecular resolution that new long-read RNA sequencing technologies can provide.

We and others have shown that the differentiation of iPSCs into macrophages produces an efferocytotic macrophage with many shared molecular features of yolk-sac derived, tissue-resident macrophages^7–9^. Here we show the molecular complexity of this process, in which alternative transcription start site (TSS) usage and isoform switching seem to play an important role in the plasticity of the iPSCs and the differentiation of myeloid progenitors and macrophages. Additionally, we observed an increase in the active TSS repertoire in stimulated monocytes and macrophages. The newly engaged TSS suggest that different isoforms of key innate immune pathway members support differences in myeloid maturation in response to different pathogens. These data open a window on hitherto uncharacterised genomic mechanisms underlying monocyte maturation and early macrophage differentiation, demonstrating the capacity for molecular expansion within the constraints of hard-wired innate gene networks.

## Results

### Section 1. Multi-omics reveals thousands of new molecules associated with iPSC and myeloid biology

To study early myeloid differentiation, cells were collected at the pluripotent, myeloid progenitor, and terminally differentiated macrophage stages (Figure 1A). These cells were examined using three different techniques (See Materials and methods): Cap analysis of gene expression (CAGE) used to assess promoter/enhancer usage; Cap-trap full-length cDNA sequencing (CFC-seq) to study isoform expression; and mass spectrometry (MS) to confirm translation of predicted open reading frames (ORF) into proteins. Despite the differences in sensitivity and molecular modality, each technique separated myeloid cells and iPSCs along the first principal component, with the second component that separated adherent cells from cells in suspension, and a smaller third component that represented differences between replicates (Figure 1B). Genes discriminating between cell types include hallmark markers of pluripotency and myeloid biology (Figure 1D, Supplementary Table 1). While data derived from FANTOM5 genome tracks demonstrate alternate transcriptional start sites at key macrophage genes (such as REL-A illustrated in Figure 1C), the CFC-seq generated here mapped these to alternate gene products and elaborated on the antisense transcripts arising from the same locus. The majority of isoforms detected by long read sequencing were supported by a CAGE peak that overlapped the transcript start site (TSS) or mapped within 100 bp upstream (Figure 1E). CFC-seq detected the largest number of expressed genes (23,625 genes called by Bambu^10^) (Figure 1F, Supplementary Table 2). 73% of these genes were associated with CAGE clusters within 500 bp, and within this group of CAGE and CFC-Seq supported genes, 54% had supporting peptides in the corresponding MS/MS datasets (Figure 1F).

**Figure 1.**
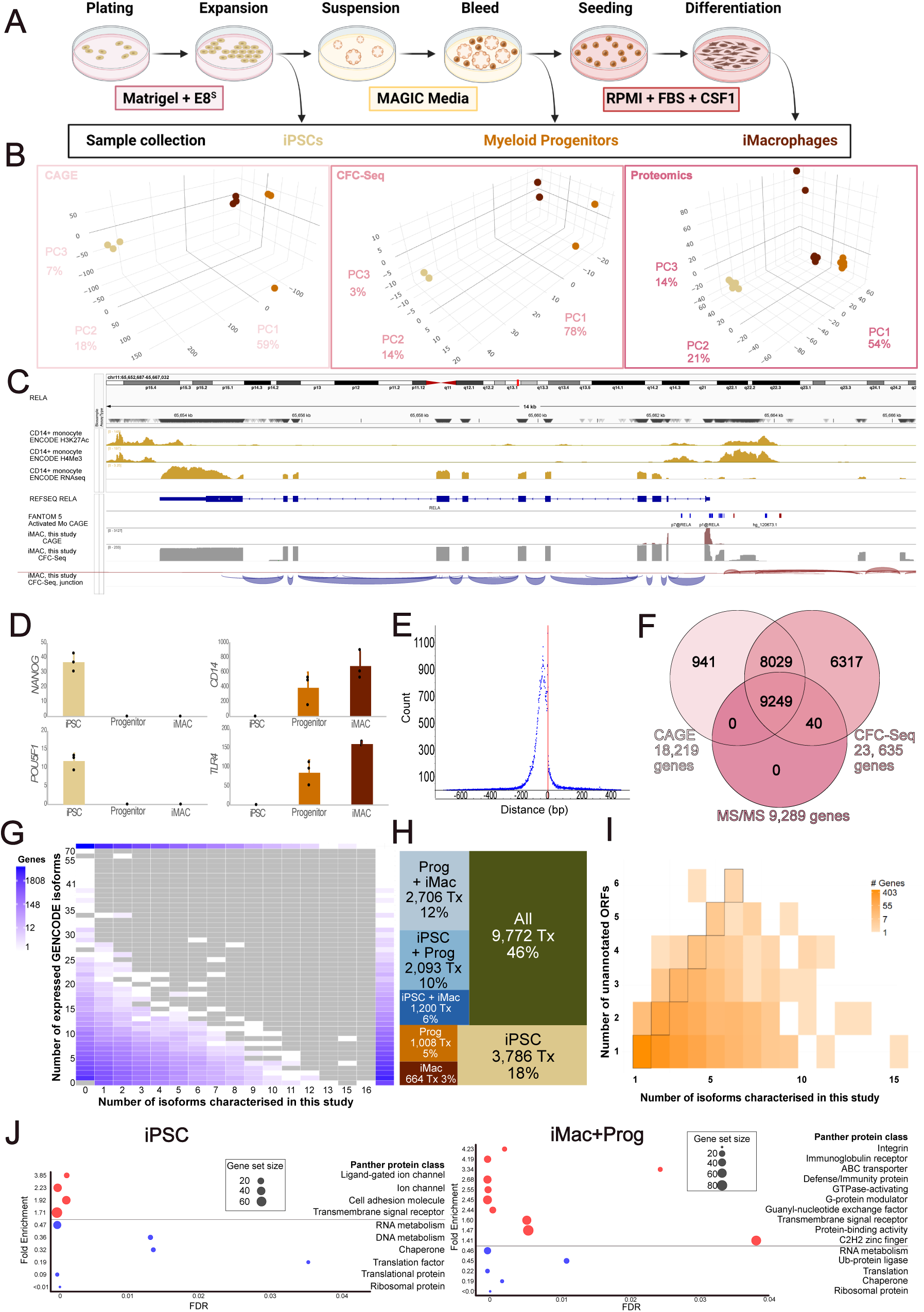
**Multi-omic approach to annotation of novel molecular features during iPSC-myeloid differentiation**. 1A) Schematic of cell models, illustrating the three cell types profiled in this study: pluripotent stem cell (iPSC), myeloid progenitor (Prog) and macrophage (iMac). Created in BioRender. Berrocal, M. (2025) https://BioRender.com/e81k263 B) PCA of major datasets (L to R) CAGE, CFC-seq and MS proteomics. C) Integrated Genome Viewer (IGV v2.19) snapshot of the *RELA* locus showing combined ENCODE CD14+ primary monocyte active promoter marks, FANTOM5 activated CD14+ monocyte CAGE track^11^, FANTOM6 iMAC CAGE and CFC-seq tracks (this study). D) Histograms summarising mean and STD of CAGE expression for cell-type markers *NANOG, POU5F1, CD14* and *TLR4*. CAGE tags are summed at gene level, biological replicates are shown as dots. E) Overlap between CAGE and CFC-seq. X-axis shows the distribution of distances between CFC-seq start positions and start position of filtered CAGE tags; y-axis shows count of events observed. The red vertical line indicates overlap at position 0 relative to CFC-seq TSS. F) Venn diagram showing numbers of genes [GENCODE v39^12^] passing library filters for each dataset and correspondence between data modalities. G) Summary of transcript diversity from all genes passing expression filters in CFC-seq and supported by tags in the CAGE libraries. The x-axis indicates the number of isoforms per gene exclusively described in this study; y-axis indicates the number of annotated isoforms that pass filtering in this dataset. Missing values in axes contain no data points. Colour represents the number of genes (log 10 scale) that match each combination of isoforms expressed in this dataset. H) Treemap summarising cell-type expression of previously uncharacterised isoforms (Tx = transcript). I) Plot showing the number of genes expressing isoforms discovered in this study (x-axis) that are predicted to contain a previously uncharacterised ORF (y-axis). Colour represents the number of genes (log 10 scale). J) Panther DB^13^ GSEA analysis of genes with at least one newly described transcript isoform, showing main protein groups found in iPSCs (left) and myeloid cells (progenitors and macrophages, right). The y-axis indicates the fold enrichment and x-axis indicates the false discovery rate (FDR). Colour indicates whether fold enrichment is overrepresented (red) or underrepresented (blue) compared to the background set, which consisted of all expressed genes that passed filtering (CFC-seq supported by CAGE and expression above threshold in at least one of the three cell types). Dot size represents number of genes from each group found in the dataset.

Using a high-confidence set of CFC-seq isoforms supported by CAGE tags, we identified 21,219 previously uncharacterised transcripts annotated to 9,277 genes (over 52% of all genes passing these filters) (Figure 1G, Supplementary Table 1). Of these, 1,105 transcripts (5.2%) mapped to genes that were not annotated in GENCODE v39, many sitting antisense to known genes (Illustrated in Fig 1C, Expanded in Section 2). Most of the uncharacterised 21,219 transcripts were expressed in all three cell types investigated (46%), followed by subsets restricted to iPSCs (18%) and myeloid cells (12%) (Figure 1H, Supplementary Table 1). This indicates that there is potential to expand the catalogues of gene products associated with pluripotency or the differentiation to myeloid cells. We predict that many of these transcripts contain ORFs that have not been previously annotated, expanding the proteome output of 1,759 genes (19% of genes to which we are adding new transcripts) (Figure 1I, Supplementary Table 3). Since alterations to the ORF may change the features or even the function of a proteoform, we examined wether different cell types are expressing alternative isoforms in different gene sets. Gene ontology (GO) analysis revealed genes with uncharacterised transcripts in myeloid cells are related to immune responses (including GTP regulated molecules like cytokines) or integrins which are involved in adhesion, survivial, proliferation and immunotolerance; on the other hand in iPSCs we mostly observed genes associated with tight junctions and ion exchange (Figure 1J, Supplementary Table 4).

### Section 2. Annotation of uncharacterised genes shows potential for discovery of myeloid secreted factors

Over 1,100 isoforms identified in this study map to 506 genes that were not present in GENCODE v39 annotations. The majority of these genes sit antisense to an annotated gene (overlapping, intronic or divergent) (Figure 2A,B). Of the 506 genes, 45 did not overlap with the domains of other known genes (intergenic) (Figure 2A,B Supplementary Table 5). Most of these newly described genes expressed multiple isoforms and most alternate isoforms shared a TSS (Figure 2B). While many of these newly characterised genes were uniformly expressed across cell types, some showed differential isoform expression between cell types (Figure 2B). Over 90% of these uncharacterised genes contain ORFs longer than 30 amino acids, suggesting they may have protein-coding potential (Figure 2B). About half of these genes had some form of annotation in the most recent genome version [GENCODE v47^12^], and a proportion were previously predicted by RefSeq GRCh38^14^ (XR_ accessions) but shown as full-length transcripts here for the first time (Figure 2B). Over 100 genes discovered in this study had not previously been reported by any of these sources (Figure 2B, Supplementary Table 5). Most isoforms annotated to uncharacterised genes were found in iPSCs (42%), followed by all cell types (18%) and myeloid cells (18%) (Figure 2C). The ORFs of newly described genes in iPSCs were enriched in proteins that contained transmembrane, cytosolic and extracellular domains, whilst myeloid cells had a majority of predictions matching extracellular proteins with a signal peptide (Figure 2D, Supplementary Table 6). This suggests that these uncharacterised genes potentially code for proteoforms that are associated with the biological functions that iPSCs and myeloid cells perform.

**Figure 2.**
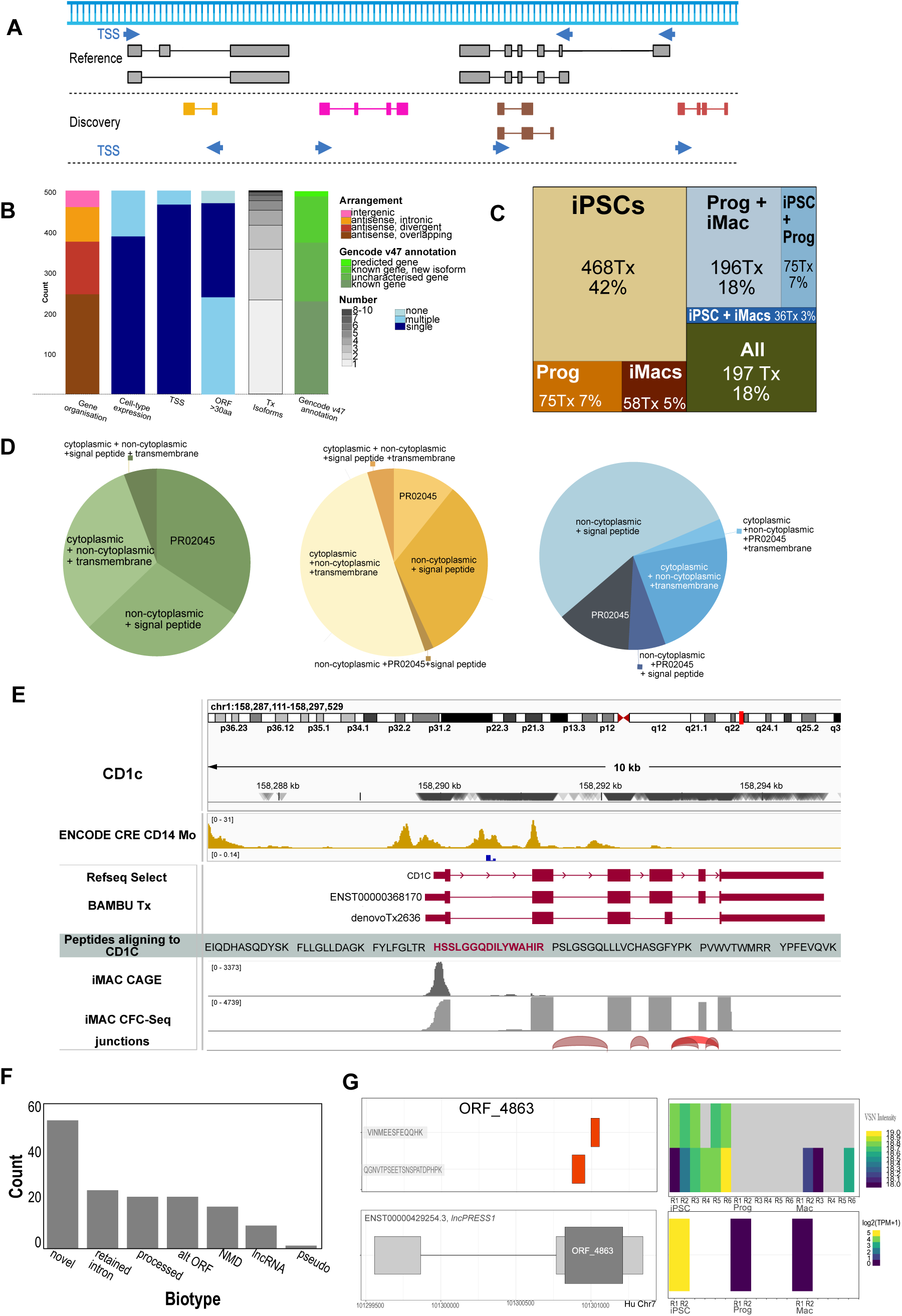
Discovery and annotation of open reading frames associated with newly described or previously annotated noncoding genes. A) Schematic illustrating GENCODE v39 reference transcripts (top) and discovery annotations (bottom) used to describe newly characterised genes. Transcriptional Start Sites for each gene indicated by blue arrows. Annotations from left to right: antisense intronic (gold); intergenic (pink); antisense overlapping (dark brown); antisense divergent (red). Created in BioRender. Berrocal, M. (2025) https://BioRender.com/y34t340 B) Descriptions of the 506 gene discovery set using categories (x-axis): <u>Gene organisation</u> relative to nearest reference gene (colour code as per (A)); <u>Cell type expression pattern</u> pale blue – all isoforms annotated to a gene are expressed in the same cell type(s); dark blue-multiple isoforms annotated to a gene are expressed in different combinations of cell types; <u>Number of TSS annotated per gene</u> dark blue - single TSS; light blue -multiple TSS; <u>Genes with predicted ORFs (>30 aa)</u> grey-blue - no ORF predicted; light blue - multiple ORF predicted; dark blue - single ORF predicted. <u>Number of isoforms</u> <u>identified per gene</u> grey scale- palest indicates single isoform, darkest grey indicates 8-10 isoforms; <u>Annotations compared to GENCODE v47</u> dark green - match to v47; forest green - new isoforms found for gene present in GENCODE v47 that was not annotated by v39; bright green - Gene predicted in RefSeq; fluorescent green – gene not previously annotated. C) Treemap showing cell type expression of isoforms annotated to previously unannotated genes (Tx = transcript; % relative to the discovery set). D). Top 5 predicted (Interpro (Blum et al., 2025)) protein motif combinations (excluding disordered domains) from 478 coding genes (988 coding Tx) discovered in this study. Pie charts coloured by cell-type distributions of predicted Tx from left to right Green (expressed in all cell types); wheat-gold (expressed in iPSC) and blue (expressed in myeloid cells). PR02045 is a highly conserved N-terminal domain. E) IGV view of CD1C, an example of a newly described proteoform of a known gene. From top to bottom: Chromosome view and 10kb window spanning CD1C locus. ENCODE composite chromatin regulatory regions identified in primary human monocytes. Reference sequence for CD1C showing intron-exon structure. Bambu-transcripts identified from CFC-seq. Peptide annotations (not an IGV track) overlayed on IGV output were derived from MS analysis. Highlighted peptide is unique to denovoTx2636. IGV tracks for CAGE data from iMAC and CFC reads plus junction predictions illustrate data contributing to BAMBU Tx annotations. F) Previously unannotated ORF validated with uniquely mapping peptides. Novel – biotype not previously described (in GENCODE v39). Retained intron; processed; alternate ORF in known gene; nonsense mediated decay (NMD); lncRNA; transcribed pseudogene. G) An example of two uniquely mapped peptides to ORF 4863 (top left) along a transcript from the lncPRESS1 gene (bottom left). Genomic location of two uniquely mapped peptides to a previously unannotated ORF (4863) with amino acid sequences indicated (top left panel) encoded by a transcript from the lncPRESS1 gene (bottom left panel). Heatmaps show expression levels across peptide samples (MS, top right panel) and long-read transcript (CFC-seq, bottom right panel), with highest expression in both modalities observed in iPSC libraries.

### Section 2.5. Mass Spectrometry confirms translation of novel isoforms

Mass spectrometry detected over 79,000 different peptides that mapped to 9,289 genes (Supplementary Figure 1). 985 transcripts with putative novel open reading frames were identified with CAGE and CFC-Seq in this study, mapped to 470 genes and their translated products were confirmed by MS (Supplementary Tables 3 and 7). This includes newly described proteoforms of CD163, a myeloid transmembrane protein implicated in tissue regeneration and inflammation^15^ and CD1c, which allows antigen presentation to T cells^16^ (Figure 2E). Although relatively few peptides (150 in total) mapped uniquely to a single isoform, in part because proteoforms may vary by a small number of amino acids, most newly described ORF were supported by multiple peptides. We identified 35 transcripts where we had high confidence that we had identified a new proteoform of known genes based on this uniqueness, including CD1c (Figure 2E, Supplementary Table 7B). Additionally, peptides uniquely mapped to six genes with transcripts annotated as ‘lncRNA’ (Figure 2F; Supplementary Table 7), including LncPRESS1 (Figure 2G), a target of p53 that is involved in maintaining pluripotency by preventing differentiation^17^. Hence, our high confidence multi-omic approach allows us to not only to confirm the existence of novel proteoforms, but also identify proteins like those arising from lncPRESS1 that could be involved in important processes like pluripotency/differentiation.

### Section 3. Alternative TSS usage is a key driver of isoform switching

Alternate mRNA isoforms were predominantly generated because of changes in the initiating or terminating exons, which accounted on its own for 64% of isoforms compared to the most similar reference sequence (in this case, most similar Ensembl transcript, Fig 3A,B, Supplementary Table 8). Limitations from the techniques used (5’ biased) and the difficulties determining alternative termination sites (Supplementary Figure 2), and the importance of TSS selection in isoform switching and function led us to focus on alternative use of initial exons. The use of an alternate 5’ exon was also predictive of internal splicing events, for example 84% of cassette exon inclusion were preceded by an alternate 5’ exon. To examine transcription initiation more closely, we reviewed the distribution of 57,747 CAGE tags (Supplementary Table 9A) between the three sample groups. Most were observed in all three cell types (∼80%, Figure 3C), with high concordance between myeloid progenitors and macrophages. Distal transcribed cis-regulatory elements (CRE), potentially enhancers, represented 10% of the CAGE tags mapping to SCAFE-predicted CRE, and the proportions of distal:proximal CRE predicted by SCAFE were equivalent between cell types (Figure 3D, Supplementary Tables 9B to 9G). Nevertheless, transcription factor (TF) motif analysis associated with CAGE tags mapping to enhancers (distal CRE) or promoters (proximal CRE) predicted a high degree of transcriptional plasticity in iPSC enhancers, with 25 transcription factor binding motifs (TFBMs) predicted to uniquely map to these regions, and a further 20 TFBMs were shared with promoters, consistently largely of known pluripotency-associated factors such as POU5F1 and SOX family members (Figure 3E, Supplementary Tables 9H to 9V). In contrast myeloid-associated enhancers appeared to be relatively fixed in the homeostatic state, with just 2 TFBM uniquely associated with these regions, whereas the proximal promoters of myeloid cells were associated with 53 unique TFBMs, plus 9 TFBMs shared with myeloid enhancers, including many known regulators of macrophage maturation and activation, including PU.1, SPIB, MEF2D, RUNX and AP-1 (Supplementary Tables 9H to 9V). This suggests that chromatin remodelling of enhancers is important for pluripotent stem cells, consistent with the idea that these cells are primed for differentiation, while committed myeloid progenitors and macrophages are poised for rapid transcriptional responses, including choice of alternate promoters within key macrophage genes.

**Figure 3.**
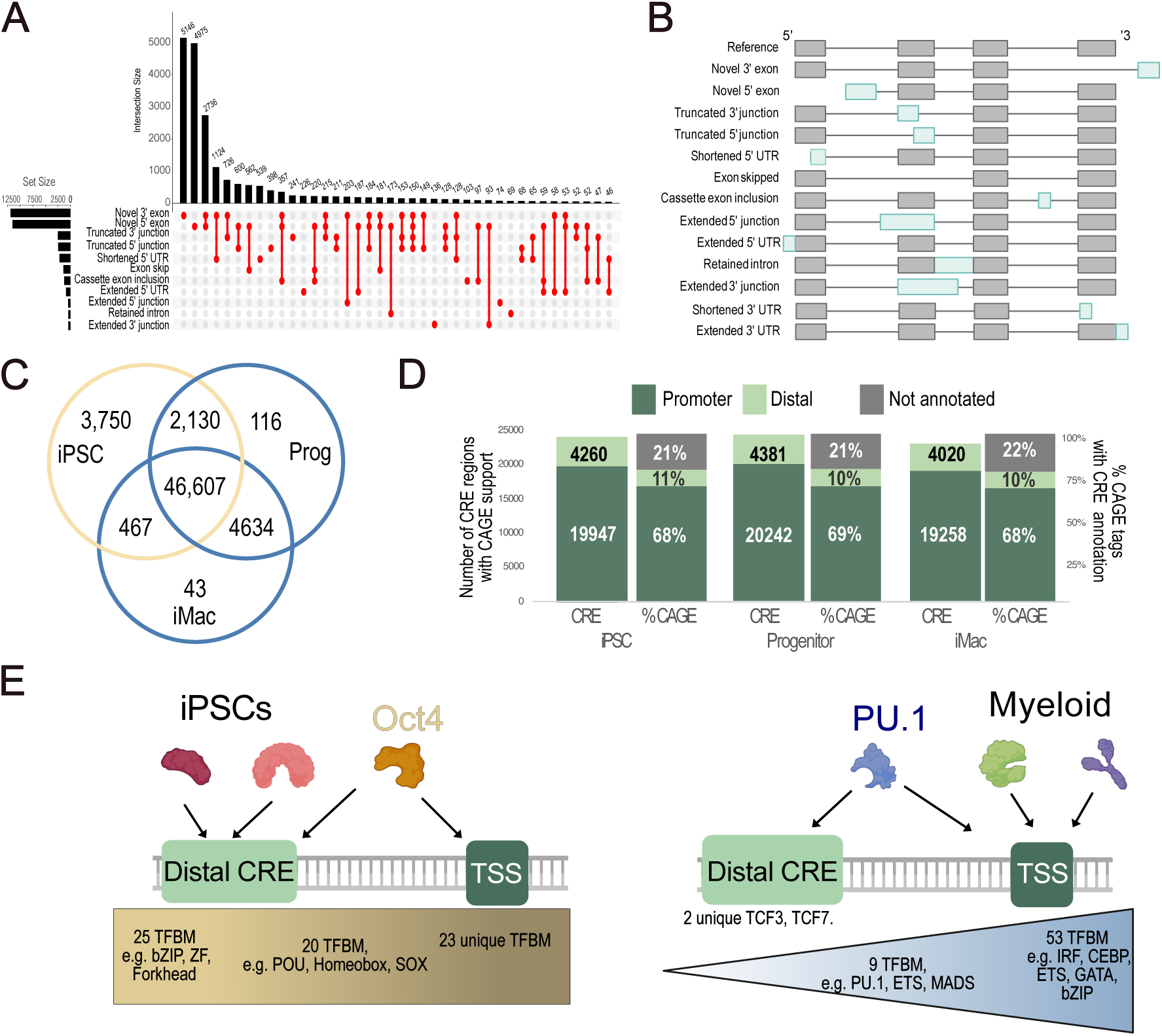
Alternative isoforms in myeloid differentiation arise from alternative TSS with changing motifs. A) Upset plot shows occurrences of alternate transcription events observed in CFC-seq data. Set size indicates number of individual transcription event types, intersection size indicates number of combined events. B) Schematic illustrates types of alternative isoforms measured in panel (A, changes in 3’ UTR not included). Reference transcript shown in grey. Pale blue sections highlight changing features. C) Venn diagram showing number of CAGE peaks expressed (sum CPM > 0) in each cell type. Pale yellow shows iPSC CAGE tags, and blue shows both myeloid cell types. D) Stacked bar plot showing the number of CRE regions supported by CAGE data (left) and % of CAGE tags mapping to CRE regions (right) for each cell type. Promoters/ proximal CRE (dark green) and enhancers/ distal-CRE (light green) were called using SCAFE (Moody et al., 2022), and those that did not map to CRE elements are shown in grey. E) Results of TF motifs significantly enriched (q < 0.01) in iPSC (left panel) and myeloid (right panel) CAGE peak regions that intersected CRE regions. Representative motifs enriched in distal regions, proximal regions and both regions are shown in the left, right and centre of each panel under the DNA schematic. Created in BioRender. Berrocal, M. (2025) https://BioRender.com/y75a785.

### Section 4. Alternative TSS usage in myeloid growth factors leads to changes in growth factor position, processing and signaling

Most ORFs we predicted for uncharacterised myeloid genes match the profile of a secreted molecule. Since these macrophages are unstimulated and resemble yolk-sac tissue-resident macrophages (known to be reparative/morphogenic)^2^, we wanted to investigate the expression profiles of growth factors. We extracted a list of all genes that match the GO term “Growth factor” from QuickGO^18^ and then studied the expression of all growth factors and their mediators in our dataset (Supplementary Figure 3). This included 62 undescribed growth factor mRNA isoforms of which 23 were predominantly myeloid (Figure 4A). Based on our results showing the importance of alternative TSS/5’ exon, we explored if this was also an important mechanism in this growth factor subset. We found that in myeloid growth factors, the use of alternative TSSs was accompanied by changes in the ORF that affected essential functional domains of these growth factors (Figure 4B). These isoforms are generally shorter, and lack positional or binding domains typical of the canonical propeptides expressed by stromal cells, suggesting the myeloid isoforms might be more bioavailable as these isoforms override control mechanisms like proteolytic release (Figure 4C). We show a previously reported case that corresponds to *IGF1 class II* ^19^, a case we had recently described with *NRG1 class VII isoforms*^20^ and a new case within this study with *TGF-β1 (tentatively termed class II)*. Alternative TSSs usage leading the expression of several uncharacterised growth factor isoforms expresed by myeloid cells were also observed for the mitogen Pleiotrophin (*PTN* denovoTx15581).

**Figure 4.**
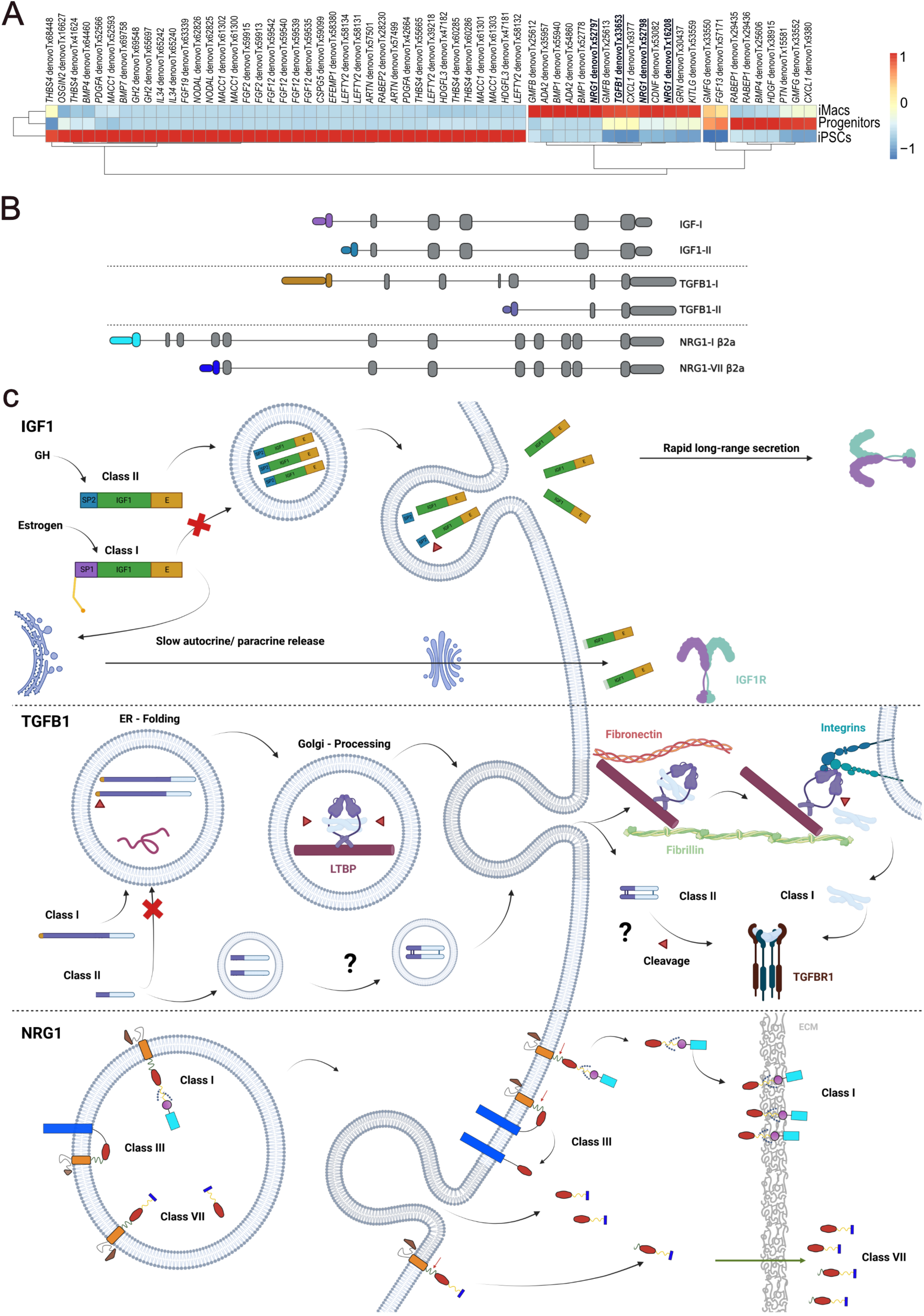
Alternate promoter use drives alternative mRNA isoforms predicted to alter location, processing and signalling of encoded growth factors. A) Heatmap showing mean expression (CFC-seq) of isoforms annotated to growth factor genes. Columns show expression of denovo predicted isoforms mapping to these genes, colour-scale indicates column normalised z-score. Underlined isoforms highlight examples drawn in schematic (B and C). See also Supplementary Figure 3 for heatmap of all growth factor transcripts. B) Transcript structure of growth factor isoforms described here, compared with representative canonical isoforms in their corresponding loci. Created in BioRender. Berrocal, M. (2025) https://BioRender.com/fhjuvnl C) Schematic of predicted consequences of myeloid-derived growth factor isoforms showing IGF-1^19^; NRG1^20^ and TGFB1 isoform (discovered this study). The common predicted features are N-terminal changes that have been shown (IGF1) or are predicted (TGFB1, NRG1) to alter pro-peptide trafficking, processing and bioavailability. Created in BioRender. Berrocal, M. (2025) https://BioRender.com/v80z072.

### Section 5. Immune activation leads to an increase in alternative TSS use

The FANTOM5 promoter atlas contains a broad repertoire of primary unstimulated and activated myeloid cells including LPS-stimulated monocyte-derived macrophages (0 to 48 hr time course)^21^, and monocytes activated with a panel of stimuli^11^. The majority (>80%) of gene-annotated TSS that were expressed at baseline (ie. in untreated cells or across both non-activated and activated cells) were supported by long read transcripts (within 50bp) from our iPSC-macrophage differentiation series (Figure 5A, Supplementary Tables 10A and 10B), indicating that our iPSC-macrophage model recapitulates the primary cell state. Activation was a major driver of TSS-level variation across the myeloid promoterome (Supplementary Figure 4A), and we found that although monocytes and macrophages expressed approximately 70% of their gene-annotated TSS in both non-activated and activated states, 25 to 28% of gene-annotated TSSs remained specific to the activated state of each cell type (Supplementary Figure 4B, Supplementary Tables 10A to 10D). Although we did not profile pathogen-stimulated iPSC-macrophages using CFC-seq, we did find long-read support (within 50bp) for over 67% of monocyte activation-induced TSS and 77% of macrophage activation-induced TSS via a combination of our long-read sequencing and GENCODE 39 transcript annotations (Supplementary Figure 4C, Supplementary Tables 10C and 10D).

**Figure 5.**
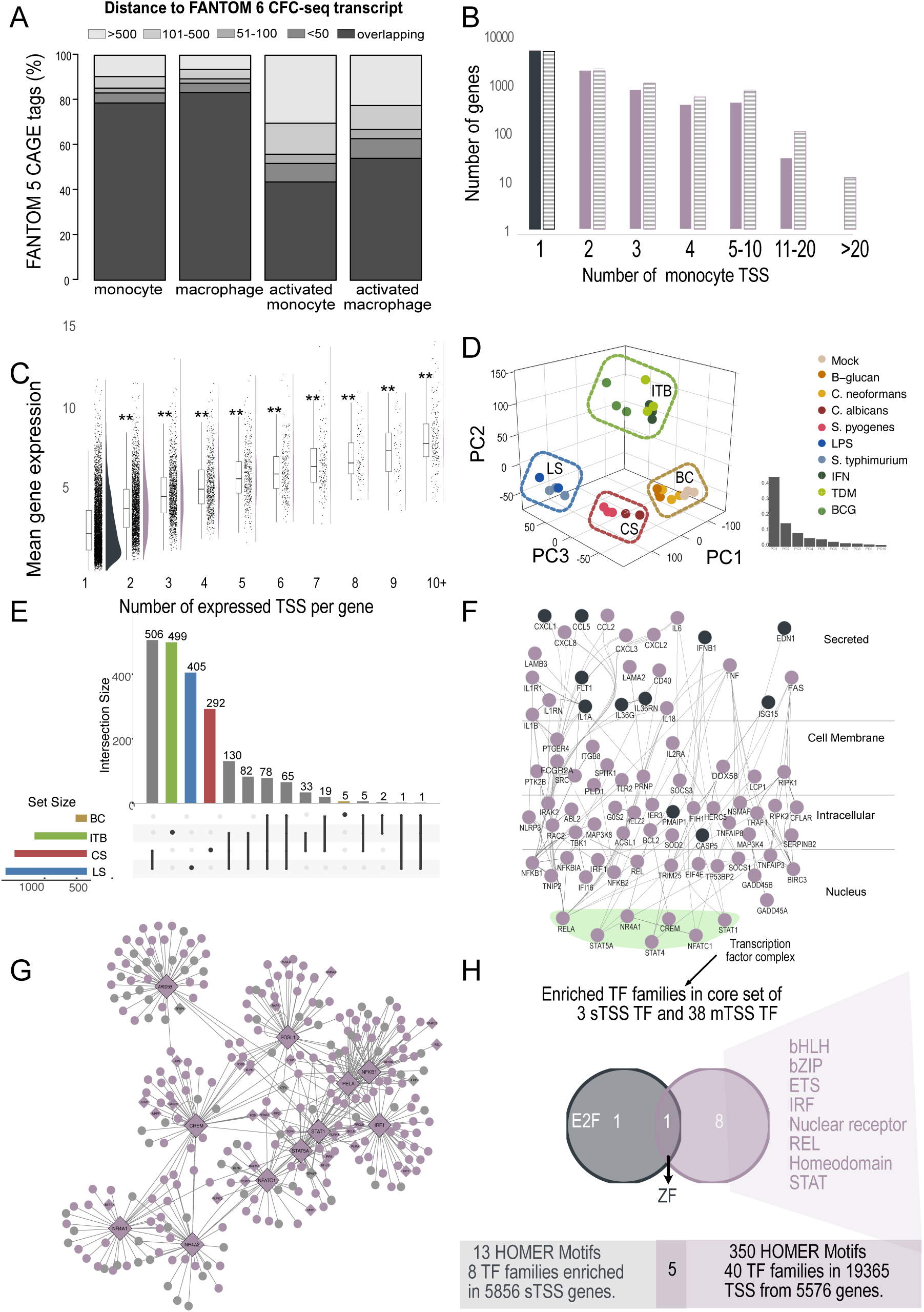
Activation of primary monocytes and macrophages leads to alternative promoter usage. A) Stacked bar plot showing distance between FANTOM5 gene-annotated CAGE TSSs and the nearest FANTOM6 CFC-seq transcript. X-axis: CAGE tags (See Supplementary Figure 4B for filtering criteria) expressed in monocytes or macrophages, or CAGE tags expressed only in the activated monocyte or macrophage libraries. Y-axis: proportion of each CAGE library found within a set distance to the nearest CFC-seq transcript. Dark grey bars indicate FANTOM5 CAGE tags overlapping the FANTOM6 CFC-seq transcript start; sequentially lighter grey bars indicate bins at <50bp, 51-100bp, 101-500bp or >500bp bins. B) Bar chart showing the number of TSSs (X-axis) used by genes expressed in primary monocytes (Y-axis, ^-^Log 10 scale). Solid bars indicate PBS-treated monocytes (n = 9995 genes, 18056 gene-annotated TSS), shaded bars indicate activated monocytes (shaded bars: n = 11283 genes, 24680 gene-annotated TSS). Grey: single TSS genes. Purple: multi-TSS genes. C) Box-whisker plot of mean gene expression for single or multi-TSS genes in primary monocytes. X-axis indicates the number of TSS observed per gene. Y-axis indicates (Log2) mean gene expression. Box whisker (median, first and third quartiles) summarises distribution of summed TSS values for individual genes, which are also plotted as discrete data points (Scatter plots) and as density distributions (grey: single TSS genes, purple: multi-TSS genes). ** indicates significant difference between sTSS and mTSS genes (Welch T-test, p<0.001). D) Analysis of monocyte activation in FANTOM5 CAGE libraries. PCA on all TSSs expressed > 1TPM in all donors for any condition. Coloured boxes highlight sample clustering subsequently used in differential expression analysis: BC: Monocytes treated with B-glucan (B) or Cryptococcus neoformans (C). CS: Monocytes treated with Candida albicans (C) or Streptococcus pyogenes (S). LS: Monocytes treated with Lipopolysaccharide (L) or Salmonella typhimurium (S). ITB: Monocytes treated with Interferon gamma (I) or Interferon gamma plus Trehalose 6,6-dimycolate (TDM) (T) or Interferon gamma plus Mycobacterium bovis Bacillus Calmette-Guérin (BCG) (B). Inset plot shows the first ten principal components (X-axis) and their contribution to dataset variance (Y-axis). E) Upset plot showing the number of significantly differentially expressed (up-regulated) gene-annotated TSSs (LIMMA treat^23^, Benjamini-Hochberg -adjusted p<0.05, minimum log(2) fold change = 1) in each sample cluster (set size) and the overlap between those differentially expressed lists (Intersection size). Coloured bars indicate lists passing significance thresholds in one cluster, grey bars in multiple groups. F) Interaction network formed from 516 genes which differentially upregulated at least one TSS across at least two pathogen clusters. Nodes: genes (grey: single TSS, purple: multi-TSS). Edges: activation, binding or inhibition, with Kappa score >= 0.8. Disconnected nodes ^22^are not shown. Cytoscape^24^ lay out – cellular component, using ClueGo^25^ plugins. G) iRegulon network showing TFs (large diamonds) identified in the common activation network (F) that were expressed from multiple TSS leading to transcripts with predicted N-terminal alterations in their ORF. Edges link TFs to their top 50 target genes (small nodes). Target genes with no TSS expressed in the monocyte activation series were removed. Purple: multi-TSS target genes. Grey: single-TSS target genes. Small diamonds: target genes which are TFs. Circles: target genes which are not TFs. iRegulon predictions were not available for the following core response TFs also predicted to express multiple protein isoforms: HIVEP1, SP110, TSC22D1 and CREB5. H) Multi-TSS genes (F) are more likely to be regulated by diverse TF families. Venn diagram showing overlap between the families of the 41 transcription factor members of the core response network (F) which were expressed from a single (grey) or multiple (purple) TSSs and had significant (q <0.05) enrichment support (at motif or family level) in the broader single- and multi-TSS gene cohort respectively. Numbers of TFs with enrichment support: sTSS: 2; mTSS: 30. Total motif enrichment in all sTSS or mTSS genes shown in lower box diagram for comparison to core network.

Isoform usage decreases as iPSC differentiated to macrophages (Figures 1 and 2), however the number of genes engaging multiple transcription start sites increased substantially in monocytes and macrophages responding to bacterial or fungal challenge (Figure 5B, Supplementary Figure 4D, Supplementary Tables 10E to 10H). Inflammatory genes engaging multiple TSS were also expressed at a higher amplitude than genes expressed from a single TSS, suggesting that this is a mechanism to promote key inflammatory outcomes in the acute phase of an infectious response (Figure 5C, Supplementary Figure 4E, Supplementary Table 10I).

To determine whether the changes we observed are a generic consequence of activation, or whether the nature of the stimulus also influences TSS dynamics, we explored TSS engagement in primary human monocytes exposed to ten different stimuli for two hours (Figure 5D). The TSS expression profiles induced by these stimuli clustered into four main groups consistent with known receptor-activated signalling pathways including TLR2-CLEC4E (*C. albicans* and *S. pyogenes*), TLR4 (*S. typhimurium* and LPS) and IFNg signalling (IFNg, *BCG* and TDM). The IFNg group (ITB) showed the highest number of uniquely induced TSSs (Figure 5E, adjusted P-value < 0.05, minimum log (2) fold change = 1), followed by the TLR4-LPS (LS) and the TLR2-CLEC4E (CS) groups. In contrast, the LS group downregulated the highest number of unique TSSs, followed by the CS and ITB groups (Supplementary Figure 4F, Supplementary Tables 10J to 10M).

We identified a core set of 516 genes which were differentially upregulated from at least one TSS, and in at least two response groups (Figure 5F, Supplementary Table 10N). Over 83% of genes in this core response network were expressed from multiple TSSs (Figure 5F) (often with disparate expression patterns (Supplementary Figures 4G and 4H) including the majority (38/41) of TFs present in the core network (Supplementary Table 10N). Based on proximity to GENCODE v39 transcripts or transcripts present in our iPSC-macrophage CFC-seq data, we predict seven core TFs were expressed as multiple proteoforms translated from different starting methionines thus potentially propagating isoform specific network outcomes depending on the type of activating stimulus. These were CREB5*, CREM*, RELA*, TSC22D1*, FOSL1, ARID5B, SP110 (* indicates transcripts for at least two proteoforms are differentially or abundantly expressed, Supplementary Figures 4I to 4L)). A further seven factors, (HIVEP1*, IRF1*, NFATC1*, NFKB1*, STAT1*, STAT5A*, NR4A1*, NR4A2) were expressed from TSSs associated with multiple GENCODE v39 transcripts likely to give rise to different proteoforms, although we could not predict which transcript/protein was expressed from CAGE data alone (Supplementary Figures 4M to 4S). iRegulon^22^ was used to identify the top 50 target genes for 11 predicted multi-protein TFs with available target gene predictions (Figure 5G). Of the predicted target genes expressed in our dataset, 72.8% were expressed from multiple transcription start sites including a further 30 target genes which were themselves TFs (Supplementary Table 10O). As each of these factors in turn have their own sets of target genes, the scope to significantly expand and fine tune the network via alternate TSS engagement at even a small number of key genes must be recognised and investigated in future studies.

Genes expressed from a single TSS were functionally enriched for ‘housekeeping’ ontologies, whereas genes enriched for functions related to immune responsiveness were predominantly expressed from multiple TSSs (Supplementary Tables 10P to 10S). These differences were reflected in the TF motifs enriched in the proximal promoters of each gene set, with single TSS genes driven by a restricted repertoire of housekeeping-associated factors, including Ronin, a TF shown to be involved in the organisation and regulation of expression from house keeping genes^26^ (Supplementary Tables 10T and 10V). In contrast, multi-TSS genes were significantly enriched for over 350 motifs across a broad range of TF families, such as NFkB/REL; STAT, IRF and bZIP which were factors present in the core response network (Figure 5F; Supplementary Tables 10U and 10W). These findings demonstrate that changes in TSS engagement may play a role in driving and shaping innate immune responsiveness.

## Discussion

Efforts to characterise the transcriptional diversity of tissue resident macrophages during early development^27^ and in adult tissues^28^ have shown great transcriptional complexity in these cells. The relative influence of “seed vs soil” or the capacity of different macrophage subsets to overlap in their specialized functions are outstanding questions in this field, and the answers may be partially found through the deep characterisation of the transcriptional networks expressed by these cells. Our in vitro approach using iPSC-derived macrophages, which models foetal yolk sac-derived myelopoiesis, suggests that alternate transcription, which mainly arises from an alternative promoter, is a feature of differentiation along the myeloid lineage. This observation is accompanied by a shift in TF motif diversity, which sits equally at enhancers and promoters in pluripotent stem cells, but shifts towards promoters as cells commit to a myeloid identity. The change in motif distribution underscores the dynamic reprogramming of transcriptional networks required during cell differentiation and the shift in the elements involved in cell plasticity at different levels or commitment.

Diversity of motifs is likely tightly linked to diverse response programs that are activated when certain stimuli are presented (pathogen epitopes, pro- or anti-inflammatory molecules, tissue-specific differentiation signals etc). We observe that many transcripts we describe here arise from genes functionally annotated as receptors or integrins, confirming the importance of microenvironmental awareness for these cells and suggesting specialisation at the receptor level for different ligands. In addition, we observe that primary monocytes respond to a variety of pathogens by activating specific sets of transcripts, which suggests their response is informed by the challenge that triggered it^29^. This diversity is also seen during differentiation, where we describe macrophage-derived growth factors that use alternative promoters potentially changing the location, processing and signalling of the proteoforms they express. Most undescribed myeloid genes shown here also fit the profile of a secreted effector, further increasing the diversity of the response repertoire. It is possible that we have previously underestimated this diversity due to several reasons: the sensitivity limitations of proteomic methods, bias towards whole cell lysates and not the secretome, or the high levels of sequence overlap preventing the use of specific antibodies at the protein level or accurate transcript assembly with short reads at the transcript level. We address this gap here and show that differentiation and specialisation as well as stimulation are accompanied by promoter switching and alternative isoform usage.

The scale of this phenomenon is considerable, even with our limited number of cell types and conservative approach: we have reported 21,219 transcripts and 506 genes that were not present in GENCODE v39. Others using similar techniques but more relaxed approaches to maximize discovery report discoveries of transcripts and genes an order of magnitude higher than we do^30^. These observations indicate we are far from a full characterisation of the features of the human genome the products that derive from it. Future work in this regard may consider characterising a wider diversity of cell types or conditions. However, any additions to the annotations must also be followed up by validation, characterisation and the interpretation of the biological significance of these products. Proteoforms are becoming recognised as an important source of functional diversity^31^, with the potential to add complexity to described molecular mechanisms or to explain pathogenic conditions. Therefore, annotation is but a necessary initial step towards functional characterisation.

In conclusion, we have evidenced that macrophage plasticity derives from molecular diversity, and that such diversity is still not fully characterised nor understood. We have shown that for myeloid cells, alternate promoter and isoform usage are leading mechanisms generating these myeloid-specific products. Lastly, we demonstrated how these products may be very important for both sensing and responding, two basic but essential features of tissue resident macrophages.

## Methods

### Computational implementation and data analysis

A complete list of software used in the analyses of these data are provided in the extended methods. Packages are referred to in the methods text here and URLS provided in Tables 1 and 2 in the extended methods file.

#### Cell lines

The use of the stem cell line PB001.1 (MCRIi001-A), obtained from the Stem Cell Core facility at the Murdoch’s Children Research Institute^32^ was in accordance with The University of Melbourne ethics committee guidelines (approval 1851831).

### Cell culture

#### Stem cell culture

PB001.1 iPSCs were cultured on growth-factor reduced Matrigel® Matrix coated petri dishes (Corning®; 356234) with Gibco™ Essential 8™ media with Essential 8™ supplement (Thermo Fisher Scientific; A1517001). Cells were cultured with daily changed fresh media. Cell culture was performed in an APT.line™ C150 (E2) CO_2_ manual incubator (BINDER; 7001–0172) in constant and stable conditions of humidity, temperature (37°C) and CO_2_ concentration in air (5%).

Cell passaging was performed routinely when cell confluency reached 70–80%. Cells were first washed with Gibco™ PBS (Thermo Fisher Scientific; 10010023) then detached using a dilution of sterile 0.5 M EDTA (Thermo Fisher Scientific; 15575020) in PBS (final concentration 0.5 mM). Cells were collected after 3 to4 min of incubation at 37°C, then pelleted through centrifugation (Heraeus Multifuge 1S-R) at 350 x g for 5 minutes at room temperature. Excess media was then eliminated, and cells were resuspended in fresh media and replated.

#### iPSC derived myeloid progenitors and macrophages

iMAC (iPSC-derived macrophage) differentiation was performed as previously described ^8,33^. Briefly: harvested cells were cultured in MAGIC media^34^. For the 3D (Embryoid body) differentiation process, PB001.1 iPSC were plated in 10 cm non-treated Petri dishes (IWAKI; 1020–100) and placed on an orbital shaker (N-Biotek orbital shaker NB-T101SRC) in a humidified incubator with 5% CO_2_ at 37 °C.

After at least 11 days of culture, cells can be observed detaching from the embryoid bodies or endothelial sheet, remaining in suspension in the media as non-adherent cells (which are characterised as myeloid progenitors). When all cells were collected and allowed to settle in a 15 mL Falcon tube (Corning®; CLS431470-500EA), embryoid bodies pelleted in the bottom, but progenitors stayed in suspension in the supernatant and could then be collected. The supernatant was centrifuged (Heraeus Multifuge 1S-R) at 400 rpm for 5 min to separate the progenitors from the media. For myeloid progenitor samples, cells collected on days 11, 13, 15 and 17 were pooled in equal concentrations.

For macrophage differentiation, progenitors collected between day 11 and 17 of the myeloid differentiation protocol were pooled and resuspended in macrophage maturation media, consisting of a 10% dilution in volume of FBS and 100 ng/mL CSF-1 (R&D Systems; 216-MC-500) in Gibco™ RPMI-1640 media (Thermo Fisher Scientific; 11875093). Cells were plated in Costar® 6-well tissue-culture treated plates (Corning®; 3516) for 4–7 days in stable incubator conditions (humid, 5% CO_2_, 37 °C), until cells showed morphological and molecular features displayed by macrophages.

### Sample collection

#### RNA collection

Total RNA was extracted using the RNeasyⓇ Plus Mini Kit (Qiagen; 74134) according to manufacturer’s instructions. RNA was collected in RNase-free water in 30 µL aliquots. RNA quality and concentration were measured using an RNA ScreenTape (Agilent Technologies; 5067–5576) in an Agilent 2200 TapeStation System (Agilent Technologies; G2964-90003).

#### Samples and library preparation for Direct cDNA CAGE (dsCAGE) and Cap-trap Full length cDNA (CFC-seq)

Total RNA for four replicates of each condition (iPSC; Hematopoietic progenitor cells; macrophages) were assessed for concentration and integrity. Samples were run on BioAnalyzer RNA Nano (Agilent Technologies, Agilent 2100 Bioanalyzer) to check for RNA degradation (RIN range 9.6 to 9.9). For each condition, 3 samples each containing more than 10 μg of RNA were separated into 2 tubes of 5 μg each (except for one sample where 2 samples of differentiated macrophages were pooled and reached only 7.6 μg, split into two tubes containing 3.8 μg of RNA each) to prepare two sets of three biological replicate libraries for each condition. The obtained RNA was processed using two different 5’-end capped RNA library preparation methods as indicated in their corresponding sections below.

#### Mass spectrometry / Proteomics

Adherent cells (iPSCs and iPSC-derived macrophages) were collected by first washing the media with cold (4 °C) Gibco™ PBS (Thermo Fisher Scientific; 10010023) 3 times and finally using a cell scraper (BIOLOGIX; Cat #: 70-1250). iPSC-derived macrophages were washed with PBS and changed to serum-free RPMI media one day before collection. Then, fresh PBS was added to the plate and cells were collected using a cell scraper (BIOLOGIX; Cat #: 70-1250) and pipetted into a 1.5 mL LoBind tube (Eppendorf; 022431081). Cells were then pelleted by spinning at 200 g for 5 minutes at 4 °C. Media was removed directly after, and cells were snap frozen in dry ice.

Floating cells (iPSC-derived myeloid progenitors) were pelleted by spinning at 200 x g for 5 minutes at 4 °C. Cells were then resuspended, washed with PBS and pelleted again (3 times). After the last spin, media was discarded, and cells were snap-frozen in dry ice.

### CAGE

#### Direct cDNA CAGE libraries (dsCAGE, full protocol manuscript under review)

Direct cDNA CAGE sequencing was performed exactly as described previously^35^. Briefly, 3.8 μg to 5 μg of RNA was reverse transcribed using a random N6+TCT primer and SuperScript III reverse transcriptase to create RNA-cDNA hybrids. The cap structure of the RNA was biotinylated and capped-RNAs were pulled-down using M-270 Streptavidin beads. Both free and cDNA-bound RNAs were degraded with RNase ONE and RNase H leaving only cDNA in the sample tube. Barcoded adapters were added in 5’- and 3’- ends of the cDNA and a second strand synthesis step was performed. The non-amplified, dual-indexed, double-stranded DNA libraries were pooled and sequenced on Illumina NextSeq2000.

#### Direct cDNA CAGE filtering and analysis

Mapping, clustering and peak calling were performed as described^35^. From the obtained files, further analysis was performed: a dataset with counts for each identified CAGE peak was generated for all 9 samples (3 replicates per cell type), filtering out reads and peaks assigned to sample S00 (unclassified reads / no sample assigned) and removing peaks mapping to scaffolds or alternative chromosomes. A reference GTF file was then built based on GENCODE v39 to annotate each of the CAGE peaks to genes (with a range of 500 bp and allowing annotation to multiple genes when overlaps were found). The data was then converted to counts per million (CPM) in each individual replicate. Data was then filtered by expression using a 0.25 CPM threshold (CAGE peak expressed over 0.25 CPM in all 3 replicates of at least one sample type). Finally, raw data for the filtered CAGE peaks was normalized using Relative Log Expression (RLE), converted to CPM and written into a final file. This resulted in a total of 57,747 CAGE peaks annotated to 17,701 genes.

#### Cap Trap-Seq cDNA library preparation for full-length sequencing

Cap-trap Full length cDNA libraries were prepared using a modified version of the CapTrap-Seq protocol^36^. No spike-ins were added in the samples. In order to capture all capped RNAs regardless of their poly-A tail status, the first step was to add a poly-A tail by adding 0.5 μl of *E. coli* Poly(A) Polymerase [5000 U/ μl] (NEB; M0276S) to 3.8 μg to 5 μg of RNA and incubate 15 min at 37 °C. Samples were then purified with 1.8x AMPure RNA Clean XP beads (Beckman Coulter; A63987) and propanol (Wako; 166-04836) and eluted in 40 μl of UltraPure DNase/RNase-Free Distilled Water (Invitrogen; 10977015). The purified poly-A tailed RNAs were then concentrated 20 min at 37 °C (final volume 5 μl). First strand synthesis was performed by adding 5.2 μl of dNTPs (ThermoFisher, 18427013) and 2.4 μl of the following priming oligo: OligodT 16VN_NB07 (<u>GAGATGTCTCGTGGGCTC</u>GG*AAGGATTCATTCCCACGGTAACAC*CTACG**TTTTTTT TTTTTTTTTVN** 3′, 100 μM, Eurofins). Note that the OligodT 16VN used contains the 16-T and VN sequence (bold) as used in Carbonell-Sala and Guigó, but a sequence complementary to the PCR primer (underlined) and an Oxford Nanopore Native Barcode sequence (italic) were also added. Here all libraries contain the Native Barcode NB07 and were sequenced separately, but using different Native Barcodes, libraries could be pooled before sequencing. The mixture (20 μl) was incubated 5 min at 65 °C and immediately cooled on ice for 1 min. The enzyme mix was prepared as follows: 8 μl of 5 × SuperScript IV buffer, 5 μl of RNaseOUT Recombinant Ribonuclease Inhibitor (Invitrogen, catalog num. 10777019), 2.5 μl of SuperScript IV Reverse Transcriptase (Invitrogen, 18090050), 2.5 μl of Induro Reverse Transcriptase (NEB, M0681S) and 2.0 μl of nuclease-free water. Twenty μl of this enzyme mix were added to each RNA/primer mix reaction in a total reaction volume of 40 μl. First-strand synthesis was performed by incubating the reaction at 55 °C for 60 min and hold at 4 °C. The resulting products were purified with 1.8x AMPure RNA Clean XP beads and resuspended in 42 μl of nuclease-free water. Cap-trap and RNA digestion were performed exactly as described above and in Carbonell-Sala and Guigó (2024), and the cDNA sample was completely dried out and resuspended in 4 μl of nuclease-free water. A double stranded linker was added in 5’ of the single stranded cDNA sample, with the following sequence:

5′ linker: contains a T7 promoter sequence (bold) and a sequence complementary to the PCR primer (underlined).

<u>ACCACCGAGATCTACACTCTTTCCC</u>**TAATACGACTCACTATAG**TACACGACGCT CTTCCGATCTNNNNNN-P

P-AGATCGGAAGAGCGTCGTGTA**CTATAGTGAGTCGTATTA**<u>GGGAAAGAGTGTAG</u> <u>ATCTCGGTGGT</u>-P

During the 5’ linker ligation, 4 μl of 5′ linker (2.5 μM, denatured for 5 min at 55 °C and put on ice for two minutes) was ligated with 16 μl of Mighty mix (DNA Ligation Kit Mighty Mix, Takara, catalogue num. 6023) to 4 μl of the sample (denatured for 5 min at 95 °C and put on ice for two minutes) and incubated 16 h at 16 °C. The 5′ linker ligation product was purified twice with 1.8 × AMPure XP beads to eliminate the non-incorporated linkers and resuspended in 42 μl of nuclease-free water. The resuspended sample (∼ 40 μl) was concentrated by drying for 45 min at 80 °C. The dried sample was resuspended with 16.25 μl of nuclease-free water. From this point, our procedure diverges greatly from the protocol by Carbonell-Sala and Guigó^36^.

The LongAmp Taq DNA Polymerase (NEB, catalog num. M0323L) PCR mix was prepared as follows: 5 μl of 5 × buffer, 0.75 μl of 10 mM dNTPs, 1.0 μl of forward primer (10 µM, Eurofins, ACCACCGAGATCTACACTCTTTCCC), 1.0 μl of reverse primer (10 μM, Eurofins, GAGATGTCTCGTGGGCTC), 1.0 μl of LongAmp Taq DNA Polymerase (2.5 U/μl). A volume of 8.8 μl of PCR mix was added to 16.25 μl of cDNA library (total volume 25 μl). The PCR cycling conditions were set to 30 s at 94 °C denaturation step, 10 cycles of 60 s at 94 °C, 60 s at 60 °C and 10 min at 65 °C amplification steps, followed by 10 min at 65 °C and hold at 4 °C. After amplification, 1 μl Exonuclease I (NEB, M0293S) was added directly into the reaction tube and incubated 30 min at 37 °C. The PCR product was purified twice with 1.8 × AMPure XP beads to eliminate any remaining PCR primers and resuspended in 42 μl of nuclease-free water. Libraries were quantified with Qubit (Thermo Fisher Scientific, Qubit 2.0 Fluorometer) and quality checked with BioAnalyzer. Using Oxford Nanopore Technologies SQK-LSK110 kit, 200 fmol of each library were loaded on different R9.4.1 flowcells and run separately on MinION MK1C.

#### Sequencing and QC

Basecalling was performed using Dorado basecaller (version: dna_r9.4.1_e8_sup@v3.3). After initial QC (see extended methods for details) data was filtered on the read sequencing quality (reads with mean qscore ≥ 10) and mapping score (MAPQ score = 60).

#### Data processing, isoform identification and annotation

Isoform discovery was performed on resulting BAM file using Bambu (v3.2.5)^10^ in “de novo” mode (annotations = NULL). To minimize bias introduced by reference annotations, which can constrain reads to known isoforms and limit novel isoform identification, a mixed approach was employed. In first pass, Bambu was run without reference annotations (annotations = NULL), generating a de novo GTF file which was subsequently overlapped with GENCODE GTF (v39) using GFFCompare (v0.12.6)^37^. Novel isoforms from GTF (excluding class codes ‘=’ and ‘c’ for known isoforms) were combined with the GENCODE GTF, forming an updated GTF to use as a reference for Bambu. In case of an overlap, GENCODE annotations prevailed over “de novo” annotations. Then, the bam files were run again on Bambu, this time using the hybrid annotations, and reads were assigned to specific isoforms within this combined annotation.

The GTF file (Extended_novomix_annotation.gtf) was edited as follows: To generate extended_novomix_annotation_v2.0.gtf, gene names for Ensembl genes were sourced from GENCODE v39 primary assembly and added to the gene_name attribute, or for BambuGenes, the Bambu gene identifier was used as the gene name. Transcripts assigned by Bambu to Ensembl gene loci, but on the opposite strand to the Ensembl gene they were assigned to had their gene_id attribute reannotated as “BambuGene_nas_[Ensembl gene id]” and the suffix – AS was added to their gene name attribute. In extended_novomix_annotation_v2.1.gtf and extended_novomix_annotation_v2.2.gtf, the nas_nomenclature was revised such that “BambuGene_nas” genes were assigned an identifier of “BambuGene_Ma_0 …” (where “…” is an integer assigned incrementally while progressing through the gene list, and “ma” indicates the transcript was manually assigned the annotation). These transcripts had their gene_name attribute removed but were instead assigned a manual_name attribute consisting of the original Ensembl gene_name suffixed with “-AS” to indicate that the transcript is antisense to that gene. We identified 27 transcripts mapping to 8 genes (Supplementary Table 11) that had been incorrectly misannotated as novel antisense genes, because they share exons with other uncharacterised transcripts correctly mapping to known genes. In these cases, gene assignments were corrected to match the overlapped gene. Finally, a list of 31 gene names mapping to 1198 gene identifiers were also removed from further analysis (5S_rRNA, 5_8S_rRNA, 7SK, GABARAPL3, HERC2P7, LINC01238, LINC02256, Metazoa_SRP, NPEPPSP1, SNORA62, SNORA63, SNORA70, SNORA71, SNORA72, SNORA73, SNORA74, SNORA75, SNORD116, SNORD30, SNORD33, SNORD39, TMSB15B, U1, U2, U3, U4, U6, U7, U8, Vault, Y_RNA). A mapping table comparing each of the annotation versions is provided in Supplementary Table 12.

#### Filtering long reads based on CAGE support and expression levels

Bed files were generated from the CAGE peak file and the CFC-seq extended annotation and compared using BEDtools^38^ (v2.26.0). Each isoform was matched to the ten CAGE tags that either overlapped or were upstream to the starting base for each isoform. Then, the distance between the most 5’ of the CAGE tags and the most 5’ base of the CFC-seq isoform was calculated. Only the match with the lowest distance was kept (if the distance was ≤ 1kb base pairs). This ensured only one line per isoform was kept when the conditions were met, but multiple isoforms could be supported by the same CAGE tag.

Next, we calculated the CPM of each transcript in each of the libraries. If the sum of CPM for a transcript within at least one of the cell types was equal or higher than 0.1, then the transcript was passed our filtering criteria (and was kept for analysis). If a transcript did not meet these criteria, it was discarded. These resulted in the transcript master table used across the study (Supplementary Table 1, Supplementary Figure 1)

### Proteogenomics

#### Sample preparation and mass spectrometry

Collected cells were lysed in 4% sodium deoxycholate containing 100 mM Tris pH 8.5 by tip-probe sonication. The lysate was heated at 95°C and centrifuged at 20,000 x *g* for 5 min at 4°C. Protein concentration was estimated with BCA and normalised to 10 μg/10 μL. Proteins were reduced with 10 mM Tris(2-carboxyethyl) phosphine and alkylated with 2-chloroacetamide and digested with was then digested with 0.2 µg of trypsin and 0.2µg of LysC overnight at 37°C. The digested was adjusted to a final concentration of 50% isopropanol and 1% trifluoracetic acid and peptides purified using inhouse made styrenedivinylbenzene-reverse phase sulfonate microcolumns. Peptides were separated on a Dionex 3500 nanoHPLC, coupled to an Orbitrap Eclipse mass spectrometer via electrospray ionisation in positive mode with 1.9 kV at 275 °C and RF set to 30%. Separation is achieved on a 50 cm × 75 µm column packed with C18AQ (1.9 µm) over 90 min at a flow rate of 300 nL/min. Peptides were eluted over a linear gradient of 3–40% Buffer B (Buffer A: 0.1% v/v formic acid; Buffer B: 80% v/v acetonitrile, 0.1% v/v FA) and the column was maintained at 50°C. The instrument was operated in data-independent acquisition (DIA) mode, with an MS1 spectrum acquired over the mass range 350–1,400 m/z (120,000 resolution, 100% automatic gain control (AGC), and 50 ms maximum injection time) followed by sequential MS/MS spectra across 13.7 *m/z* isolation windows with 1 *m/z* overlap covering the full mass range. MS/MS data will be acquired with higher-energy collisional dissociation (HCD) fragmentation (30,000 resolution, 2000% AGC, 55 ms maximum injection time, and normalized collision energy 30 eV).

#### Proteogenomics database creation

GenomeProt (https://github.com/ClarkLaboratory/GenomeProt) was used to build a custom proteogenomics database of potential open reading frames (ORFs) encoded by known and novel isoforms (Figure 6). Details of the pipeline are provided in the extended methods file.

**Figure 6.**
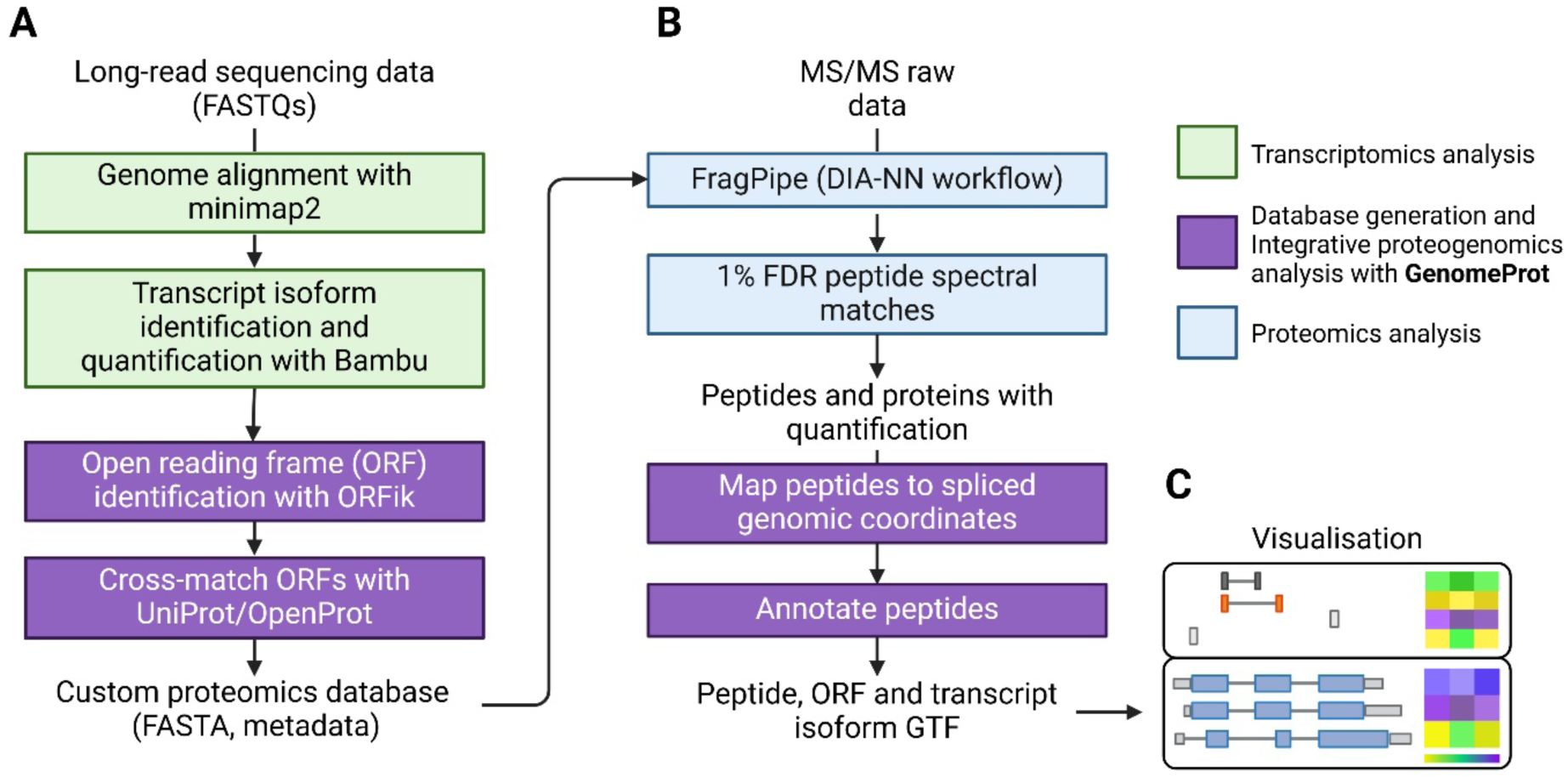
Overview of the GenomeProt workflow. A) The database generation module accepts RNA sequencing FASTQ files, BAM files or GTF annotation files from both short-read and long-read sequencing platforms. We ran Bambu with our own parameters (Methods) and subsequently entered the workflow after this step, using the GTF as input. The main output from this module is a FASTA file with known protein entries and unannotated open reading frame sequences. B) Proteomics data is processed externally using FragPipe. C) The output peptides.txt file from FragPipe is used along with the previously generated database to map peptides to spliced genomic coordinates, the isoforms that encoded them and annotate whether these mappings are unique or shared among features. This module outputs a GTF file of peptide, ORF and isoform coordinates which is used in the visualisation module to display peptide, and isoform tracks alongside quantitative heatmaps.

Reference Transcript Selection:

To systematically explore types of novel isoforms against a known gene, we developed a computational framework to identify the most appropriate reference transcript from GENCODE annotations. The reference transcript was selected based on its consistency with the novel isoform, prioritising exon structure conservation and maximising exon similarity.

#### Classification of Novel Isoform Types

Following reference gene selection, exon-level structural differences between the novel isoform and its reference gene were classified into distinct transcript modification types. Based on these exon comparisons, the following categories of isoform modifications were identified: Novel flanking exons (5’ or 3’): the boundaries of either the first (5’) or last (3’) exon of the studied isoform don’t match the reference. UTR Modifications: the 5′ UTR and 3′ UTR were compared between the novel isoform and reference gene. Extensions or truncations were classified when the exon boundaries shifted by more than 200 bp for 5′ UTR extension and 100 bp for 5′ UTR shortening, to exclude minor sequencing or annotation artifacts. In the case of 3’ UTRs, distances between transcriptional ends could not be determined, and so both extension and shortening were no longer investigated in this analysis. Cassette Exon Inclusion & Exon Skipping: a novel exon present in the isoform but absent in the reference was classified as cassette exon inclusion, whereas missing exons in the novel isoform relative to the reference were classified as exon skipping. Retained Introns: when an intronic region in the reference gene was retained within an exon of the novel isoform, the event was classified as a retained intron. Exon Boundary Shifts: if an exon boundary in the novel isoform was extended or truncated compared to the reference gene, we classified them as either Extended 5′ or 3′ junction or Truncated 5′ or 3′ junction, respectively.

#### Growth factor heatmap

A list of all genes annotated as “growth factor” within the QuickGO database^39^ was obtained, and the transcript master file (Supplementary Table 1) was filtered to only retain matching gene IDs.

#### FANTOM5 Data preparation

Annotated, normalised FANTOM5 CAGE expression data (RLE TPM hg38 fair and new phase 1 and 2 peaks) were sourced from FANTOM https://fantom.gsc.riken.jp/5/datafiles/reprocessed/hg38_v9/extra/CAGE_peaks_expression/ (last modified: 2021-05-28 18:23). CAGE-TSSs mapped to multiple Entrez or HGNC gene IDs (n = 633), were removed, leaving 209278 TSSs in the “global” FANTOM5 CAGE set. Myeloid cell type specific datasets (basophils, CD34+ progenitors, monocyte-derived dendritic cells, plasmacytoid dendritic cells, monocyte derived macrophages, monocytes (ex vivo), mast cells, Langerhans cells, AML cell lines, CML cell line, monocyte-derived macrophage LPS time course, monocyte activation series) were extracted from the global CAGE set according to the expression criteria described in Supplementary Table 10X, combined to define the “myeloid” TSS set, which consisted of 83096 CAGE-TSS mapped to 16037 gene identifiers.

#### Assessing long read CFC-seq support for FANTOM5 CAGE TSSs

FANTOM5 CAGE TSSs were mapped to the nearest FANTOM6 CFC-seq transcript expressed above threshold (expression above 0.1 CPM when adding both replicates of at least one of the cell types) (Supplementary Table 1, Supplementary Figure 1) and to a combination of FANTOM6 and GENCODE v39 transcript start site regions (extended_novomix_annotation_v2.2.gtf).

#### Assessment of TSS engagement in Monocytes and Macrophages

Genes from the FANTOM5 monocyte activation series and monocyte-derived macrophage LPS time course series were assessed as single- or multi-TSS genes in a dataset specific manner according to the number of detected TSS assigned to each Entrez (or, if unavailable, HGNC) identifier. Gene identifiers with only one detected TSS were designated as single TSS genes; genes with more than one detected TSS were classified as multi-TSS genes.

#### Locus expression in FANTOM5 samples

Locus expression levels were calculated by summing together the average expression measurements of each TSS (across all samples within each dataset).

#### Identification of differentially and abundantly expressed FANTOM5 TSSs

Differential expression analyses for all TSSs expressed above threshold in the FANTOM5 monocyte activation series were performed between each pathogen cluster (Figure 5D) and PBS-treated monocytes following the R (v4.3.1)/ Limma (v3.56.2)^23^. The mean-variance relationship was accounted for using ‘voom’, and empirical Bayes moderated t- and p-values were calculated using Limma’s ‘treat’ function with a minimum log(2) fold change of one. Significant differential expression was determined from the treat output, as an adjusted p-value cut off < 0.05. TSSs expressed above the average sample median (2.43 TPM) in all samples were classified as abundantly expressed.

#### Identification of ‘core’ gene set (FANTOM5 monocyte activation series)

The ‘core’ of differentially upregulated genes was comprised of genes which differentially upregulated TSS (excluding enhancer-associated TSS and non-gene-annotated TSS) in at least two different pathogen clusters compared to mock-treated monocytes. An interaction network was plotted in Cytoscape (v3.10.3) and the Human Protein Atlas^40^ used to inform sub cellular locations.

#### Pathway overrepresentation analysis

Pathway over-representation was assessed in each indicated FANTOM5 gene set using InnateDB (hypergeometric algorithm with Benjamini-Hochberg correction, adjusted p < 0.05)^41^.

#### CRE analysis

Regulatory elements were annotated by applying the SCAFE method^42^ to the CAGE data (each sample was analysed independently). Within each regulatory class (P/D), overlapping and book-ended CRE regions were merged in a strand-specific manner across each cell-type’s replicates. The global CAGE set for each cell type was identified by filtering detected CAGE peaks (> 0.25 CPM) (Supplementary Figure 1) to further exclude peaks without a replicate sum > 0 in the relevant cell type). CAGE peaks were mapped to CRE-P and CRE-D regions (separately) in a cell-type and strand-specific manner.

## Data availability

Raw data files are deposited in https://fantom.gsc.riken.jp/6/datafiles/

## Code availability

Code used for analysis can be found in: https://github.com/wellslab/FANTOM6

## Author contribution statement

MABR, SKB and CAW designed the study and analytical approaches.

MABR, JG, MK, DD, HK, AB, DV, KH, BLP generated and analysed data.

MABR, JG, HK, KH, CZ, MBC, BP, SKB designed code and implemented workflows for this data.

MBC, BLP, HT, PC, CAW supervised and funded the project.

MABR, JG, CZ, SKB, CAW contributed figures and wrote the paper.

## Funding Acknowledgements

MABR was funded by a University of Melbourne Research Scholarship. The project was supported by Australian National Health and Medical Research Council (NHMRC) Synergy grant APP 1186371 to CAW, NHMRC Investigator Grants GNT1196841 to M.B.C. University of Melbourne Research Flagship to M.B.C, B.L.P and C.A.W. FANTOM was supported by research grant for the RIKEN Center for Integrative Medical Sciences (IMS) from the Ministry of Education, Culture, Sports, Science and Technology (MEXT) Japan. This work was supported by the Human Technopole Foundation, a research foundation funded by the Italian Government under the Ministries of Economy & Finance, Health, and Education, University and Research.

## Other Acknowledgements

Sequencing was performed by Laboratory for Genotyping Development in RIKEN IMS and the National facilities (formerly the sequencing facility) and the Centre of Genomics in Human Technopole. We thank Nobuyuki Takeda and Teruaki Kitakura (RIKEN) for their support of the IT infrastructure for the FANTOM6 collaboration, and Emi Ito (RIKEN) for her administrative support. This is a collaborative work with FANTOM6 Consortium. We thank all consortium members for their insights and suggestions.

